# Electron tomography visualization of HIV-1 virions trapped by fusion inhibitors to host cells in infected tissues

**DOI:** 10.1101/2024.08.19.608557

**Authors:** Mark S. Ladinsky, Li Zhu, Irfan Ullah, Pradeep D. Uchil, Priti Kumar, Michael S. Kay, Pamela J. Bjorkman

## Abstract

HIV-1 delivers its genetic material to infect a cell after fusion of the viral and host cell membranes, which takes place after the viral envelope (Env) binds host receptor and co-receptor proteins. Binding of host receptor CD4 to Env results in conformational changes that allow interaction with a host co-receptor (CCR5 or CXCR4). Further conformational rearrangements result in an elongated pre-hairpin intermediate structure in which Env is anchored to the viral membrane by its transmembrane region and to the host cell membrane by its fusion peptide. Although budding virions can be readily imaged by electron tomography (ET) of HIV-1–infected tissues and cultured cells, virions that are fusing (attached to host cells via pre-hairpin intermediates) are not normally visualized, perhaps because the process of membrane fusion is too fast to capture by EM. To image virions during fusion, we used fusion inhibitors to prevent downstream conformational changes in Env that lead to membrane fusion, thereby trapping HIV-1 virions linked to target cells by prehairpin intermediates. ET of HIV-1 pseudovirions bound to CD4^+^/CCR5^+^ TZM-bl cells revealed presumptive pre-hairpin intermediates as 2-4 narrow spokes linking a virion to the cell surface. To extend these results to a more physiological setting, we used ET to image tissues and organs derived from humanized bone marrow, liver, thymus (BLT) mice infected with HIV-1 and then treated with CPT31, a high-affinity D-peptide fusion inhibitor linked to cholesterol. Trapped HIV-1 virions were found in all tissues studied (small intestine, mesenteric lymph nodes, spleen, and bone marrow), and spokes representing pre-hairpin intermediates linking trapped virions to cell surfaces were similar in structure and number to those seen in the previous pseudovirus and cultured cell ET study.

**IMPORTANCE:** Trapped and untrapped HIV-1 virions, both mature and immature, were distinguished by localizing spokes via 3D tomographic reconstructions of HIV-1 infected and fusion-inhibitor treated tissues of humanized mice. The finding of trapped HIV-1 virions in all tissues examined demonstrates a wide distribution of the CPT31 inhibitor, a desirable property for a potential therapeutic. In addition, the presence of virions trapped by spokes, particularly in vascular endothelial cells, demonstrates that fusion inhibitors can be used as markers for potential HIV-1-target cells within tissues, facilitating the mapping of HIV-1 target cells within the complex cellular milieu of infected tissues.

## INTRODUCTION

Fusion between the viral and target cell membranes is a requisite first step for HIV-1 infection of a target cell (1). The fusion process is mediated by HIV-1 envelope (Env), a trimeric glycoprotein comprising three gp120-gp41 protomers in which the gp120 subunits contain the host receptor (CD4) and co-receptor (CCR5 or CXCR4) binding sites and the gp41 subunits include the hydrophobic fusion peptide and membrane-spanning region (Fig. 1A, step i). CD4 binding to an Env trimer triggers conformational changes that include outward rotation of the three gp120 subunits and displacement of the V1V2 region to expose the binding site for co-receptor on V3 (2–5) (Fig. 1A, step ii). Upon co-receptor binding, the fusion peptide within gp41 is released to allow its insertion into the host cell membrane (Fig. 1A, steps iii and iv), after which fusion of the host and viral membranes is accomplished following formation of a six-helical bundle in which the HR1 (N-trimer) and HR2 (C-peptide) regions of gp41 form a six-helical bundle (6, 7) (Fig. 1A, steps v and vi). Preventing six-helical bundle formation with a C-peptide-based fusion inhibitor, which recognizes a site on N-trimer, blocks viral-host cell membrane fusion by trapping virions at the host cell surface in a pre-hairpin intermediate structure in which the gp41 fusion peptide is inserted into the host cell membrane and the gp41 transmembrane region remains in the viral membrane (8). Fusion inhibitors include T20 (enfuvirtide [Fuzeon]) (9), T1249, a more potent derivative of T20 (10), anti-gp41 antibodies such as D5 (11), and CPT31, a highly potent cholesterol-conjugated trimeric D-peptide (12).

**FIG 1.**
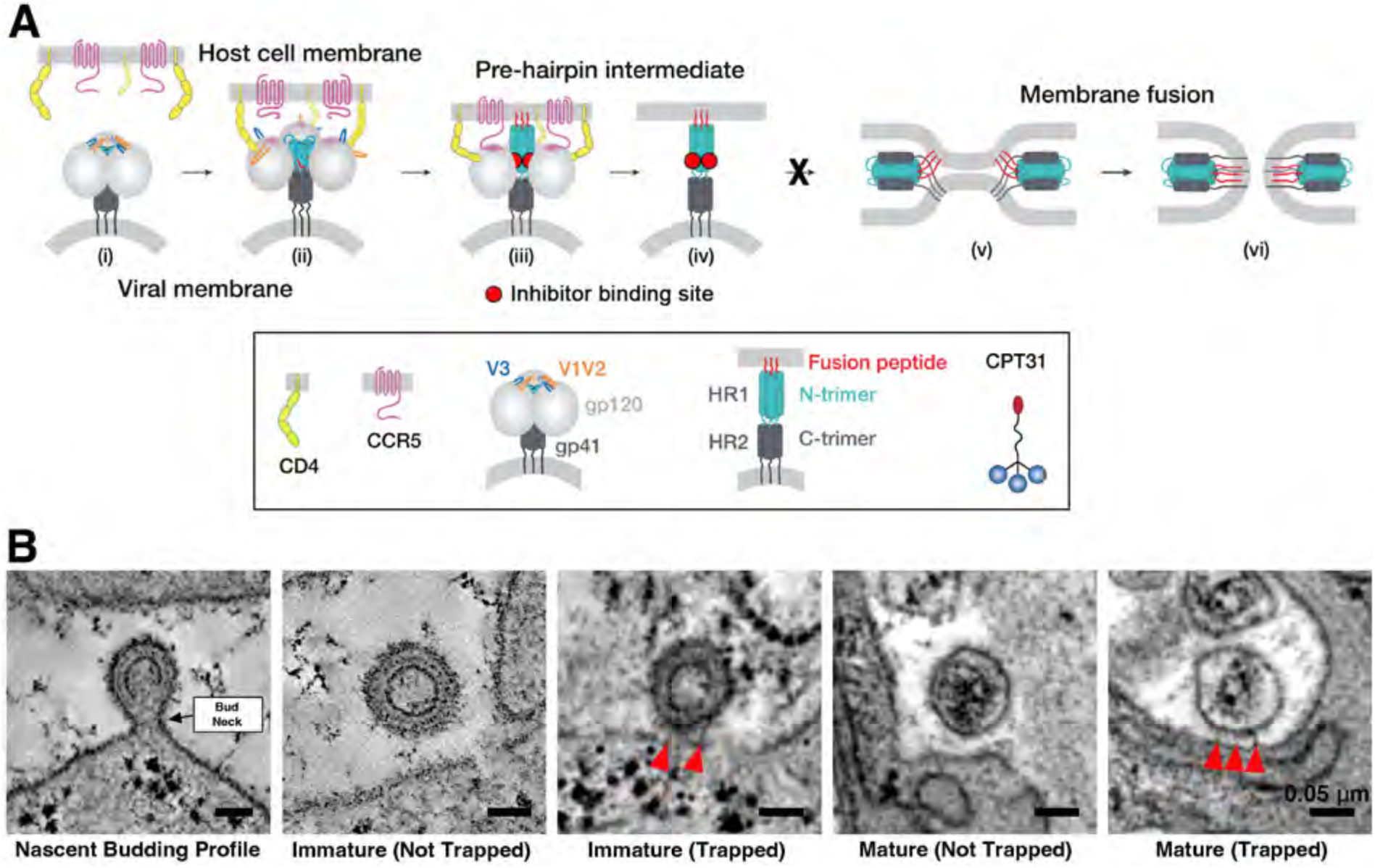
Steps involved in HIV-1 fusion and representative ET images. (A) Schematic of steps involved in Env-mediated membrane fusion. (i) Unliganded closed, prefusion HIV-1 Env trimer in which the V1V2 loops (orange) occlude the coreceptor-binding site on V3 (blue). The HIV-1 Env trimer is embedded in the viral membrane and the host receptor (CD4) and coreceptor (CCR5) are embedded in the target cell membrane. (ii) CD4-bound open HIV-1 Env trimer in which V1V2 loops are displaced to expose the coreceptor-binding site on V3. (iii) Hypothetical receptor- and coreceptor-bound open Env trimer with rearrangements of gp41 N-trimer/HR1 to form a pre-hairpin intermediate linked to the target cell membrane by the gp41 fusion peptide (red). Red circles indicate approximate binding sites for CPT31. (iv) Hypothetical pre-hairpin intermediate formed by gp41 trimer after gp120 shedding. Red circles indicate approximate binding sites for CPT31. Estimates for the maximum length of a pre-hairpin intermediate range from 370 Å (maximally extended conformation outside of helical HR1 and HR2 regions) to 450 Å (maximally extended conformation outside of a helical HR1 and an extended HR2). However, measured distances between viral and host cell membranes linked by spokes are shorter (22) (Fig. 3E), suggesting that regions outside of helical HR1 and HR2 residues are not fully extended. (v–vi) Formation of a post-fusion gp41 six-helical bundle that juxtaposes the host cell and viral membranes (step v) prior to membrane fusion (step vi). Bottom: Schematics of host receptor and co-receptor, HIV-1 Env trimer, pre-hairpin intermediate, and CPT31 fusion inhibitor. (B) Localization of inhibitor-trapped and free (non-trapped) virions in humanized BLT mouse tissues by ET. Immature and mature HIV-1 virions (both trapped and free) can be distinguished from nascent HIV-1 budding profiles. Red arrowheads; spokes. Data in panel 5 are also shown in Movie 1.

Prior to employing pre-hairpin intermediate fusion inhibitors, HIV-1 virions at stages of fusion or ingress into a target cell had not been unambiguously identified despite extensive 3D imaging by electron tomography (ET) of HIV-1–infected cultured cells (13–17) and tissues (18–21). Indeed, in our ET studies of HIV-1–infected humanized mouse tissues, we identified hundreds of budding virions and thousands of free mature and immature virions (18–21), but no examples of a virus attached to a host cell via a pre-hairpin intermediate or in the process of fusing its membrane with that of a target cell. This result can be rationalized by assuming that viral fusion is very fast compared with viral budding; thus, when infected samples are immobilized for imaging, the relatively slow process of viral budding would be more easily captured compared with the presumably faster process of fusion. We demonstrated that treatment with an HIV-1 fusion inhibitor that recognizes the exposed gp41 N-trimer after host cell receptor and coreceptor binding (Fig. 1A, steps iii and iv) allows visualization of pre-hairpin intermediate structures in trapped virions by ET: we collected >100 examples of HIV-1 pseudoviruses linked to receptor- and co-receptor-expressing TZM-bl target cells by 2-4 narrow rod-like densities (which we referred to as spokes) in inhibitor-treated samples, but none in untreated or control-treated samples (22).

Here, we extended our previous in vitro study (22) by conducting ET of tissue samples from humanized mice infected with HIV-1 prior to treatment with the fusion inhibitor CPT31 (12). This approach allowed investigation of fusion inhibitor-trapped authentic HIV-1 virions attached to target cells in tissues and organs of an actual infected animal. Similar to our previous study (22), we again found trapped virions linked to target cells by 2-4 narrow rod-like densities (spokes). Using 3D tomographic reconstructions of >100 trapped virions within humanized mouse tissues, we measured spoke dimensions and spoke spacing to create consensus models of pre-hairpin intermediate-linked virion and target cells in optimally preserved plastic-embedded ET samples. Furthermore, we demonstrated that fusion inhibitor-trapped virions can serve as markers for HIV-1 target cells within mucosal tissues of an infected organism and that the CPT31 inhibitor is distributed widely in an HIV-1–infected animal.

## RESULTS

Mouse models with humanized immune systems can be used to study aspects of HIV-1 infection in mucosal lymphoid tissues (23). Humanized bone marrow/liver/thymus (BLT) mice, which are individually created by transferring human fetal thymic and liver tissues and CD34^+^-human stem cells into immunocompromised mice, represent an optimal model of HIV-1 infection (24). Humanized BLT mice show human immune cell engraftment at mucosal sites (25, 26) and reconstitution of high levels of human lymphoid immune cells including T and B cells, monocytes, dendritic cells, and macrophages (27, 28).

For the current experiments, humanized BLT mice that had been infected with HIV-1_JRCSF_ for 3 weeks were subsequently treated with two doses of CPT31 (10 mg/kg; 2 h apart; dose chosen as described in Methods based on data in ref. (12)), a high-affinity D-peptide inhibitor linked to cholesterol (MW = 9 kDa) (12, 29). Tissue samples from HIV-1–infected/CPT31-treated mice were collected 2 h after the second inhibitor treatment and prepared for EM by high-pressure freezing and freeze-substitution prior to plastic embedding (30–33). As in our previous imaging studies (19–21), HIV-1–infected tissue samples were lightly fixed with aldehyde upon dissection to render them safe for handling outside of ABSL-3 containment (34).

To identify trapped virions, we used EM to scan the peripheries of cells within tissue sections to first locate ∼100 nm spherical objects adjacent to cell surfaces. Potential trapped virions were examined at higher magnification and at tilts of 0°, 35° and −35° to further verify their spherical nature and confirm that the objects were not part of larger extracellular projections (e.g., pseudopods), as expected for authentic HIV-1 virions. Presumptive virions were further examined using a defocus series to detect core structures that should be found inside HIV-1 virions: i.e., a cone-shaped core in mature HIV-1 and a C-shaped core in immature HIV-1 (35–37) (Fig. 1B; Movie 1). Once virions were identified, the region containing the virion(s) and adjacent cellular structures was imaged by a dual-axis tilt series for 3-D reconstruction by ET. Trapped HIV-1 virions (both mature and immature) were distinguished from untrapped virions by the presence of spokes linking them to adjacent target cell surfaces, and virions (trapped or untrapped) could be distinguished from nascent budding profiles by the presence of a bud neck on the latter (Fig. 1B).

Tissues selected for this study included small intestine (gut-associated lymphoid tissue, GALT) (Figs. 2,3), bone marrow (Fig. 4), mesenteric lymph node (Figs. 5,6), and spleen (Fig. 7). These tissues were chosen because they are rich in HIV-1 target cells that are of human origin, are known to harbor populations of HIV-1 during peak infection, and because they were examined in our previous ET studies of HIV-1 infection in humanized mice (19–21). Trapped HIV-1 virions were identified in all tissues examined from infected humanized BLT mice treated with CPT31, complementing our study of pseudovirus infection of cultured cells in the presence of fusion inhibitors (22) and confirming that virion trapping can also be observed in vivo under conditions of an infection with authentic HIV-1.

**FIG 2.**
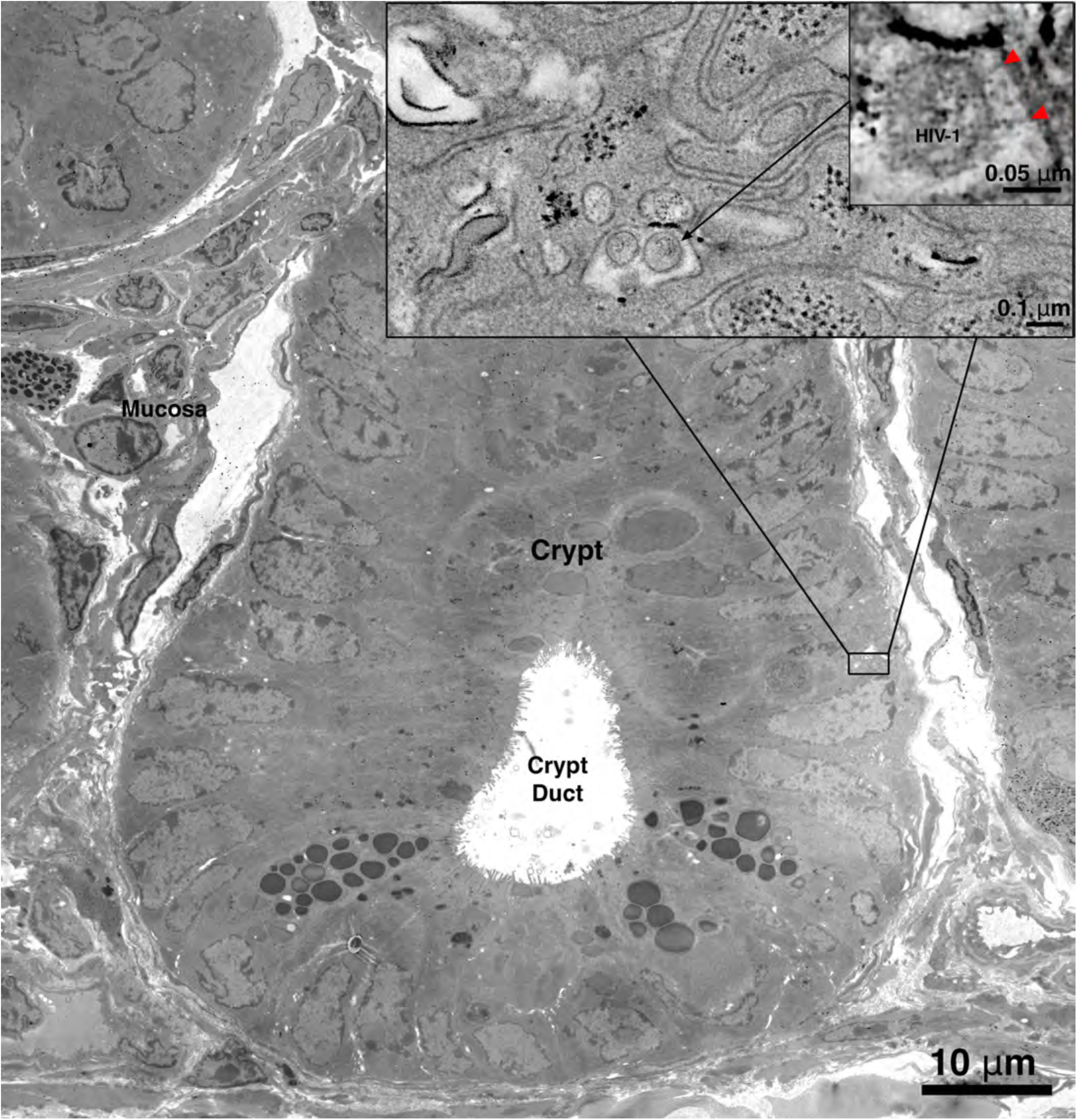
Trapped virions in GALT (small intestine). Virions were found near the peripheries of the intestinal crypts, similar to previous observations (21). Inset: ET detail of a crypt peripheral region, showing several HIV-1 virions in the intracellular space. Tomography of a single trapped virion shows attachment to the surface of an adjacent target cell by 2 narrow spokes (red arrowheads).

**FIG 3.**
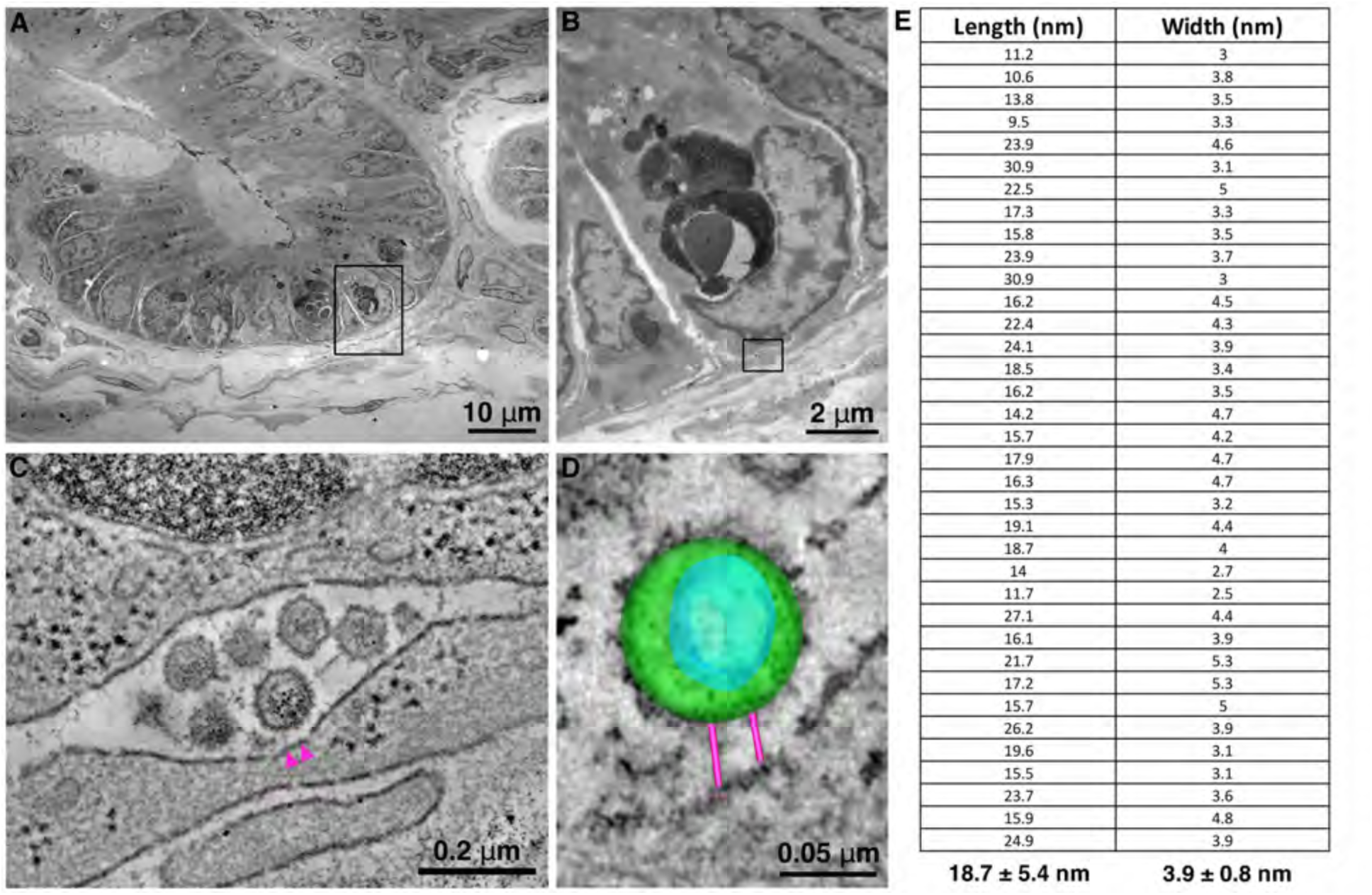
Trapped virions in GALT. (A) Montaged projection overview of the basolateral portion of a crypt in humanized BLT mouse GALT. (B) Detail of the region indicated by the rectangle in panel A, showing a pool of free virions in the intracellular space near the periphery of the crypt (rectangle). This localization of virions is similar to that seen in previous GALT ET studies (19, 21). (C) Tomographic reconstruction of the virion pool indicated by the square in panel B. Spokes (magenta arrowheads) were present on one of the six virions in the pool. (D) 3D segmentation of the inhibitor-trapped virion (green, virion; blue, viral core; magenta, spokes) from panel C. (E) Measurement (length/width) of spokes found on trapped HIV-1 virions in GALT. Trapped virions in tomographic reconstructions were digitally extracted and rotated in 3D to optimize visualization of spokes and then measured by modeling their length and width using tools in IMOD (66). The average length and width of 58 trapped virions visualized in 21 ET data sets of infected GALT were 18.7 (+/− 5.4) nm and 3.9 (+/− 0.8) nm.

**FIG 4.**
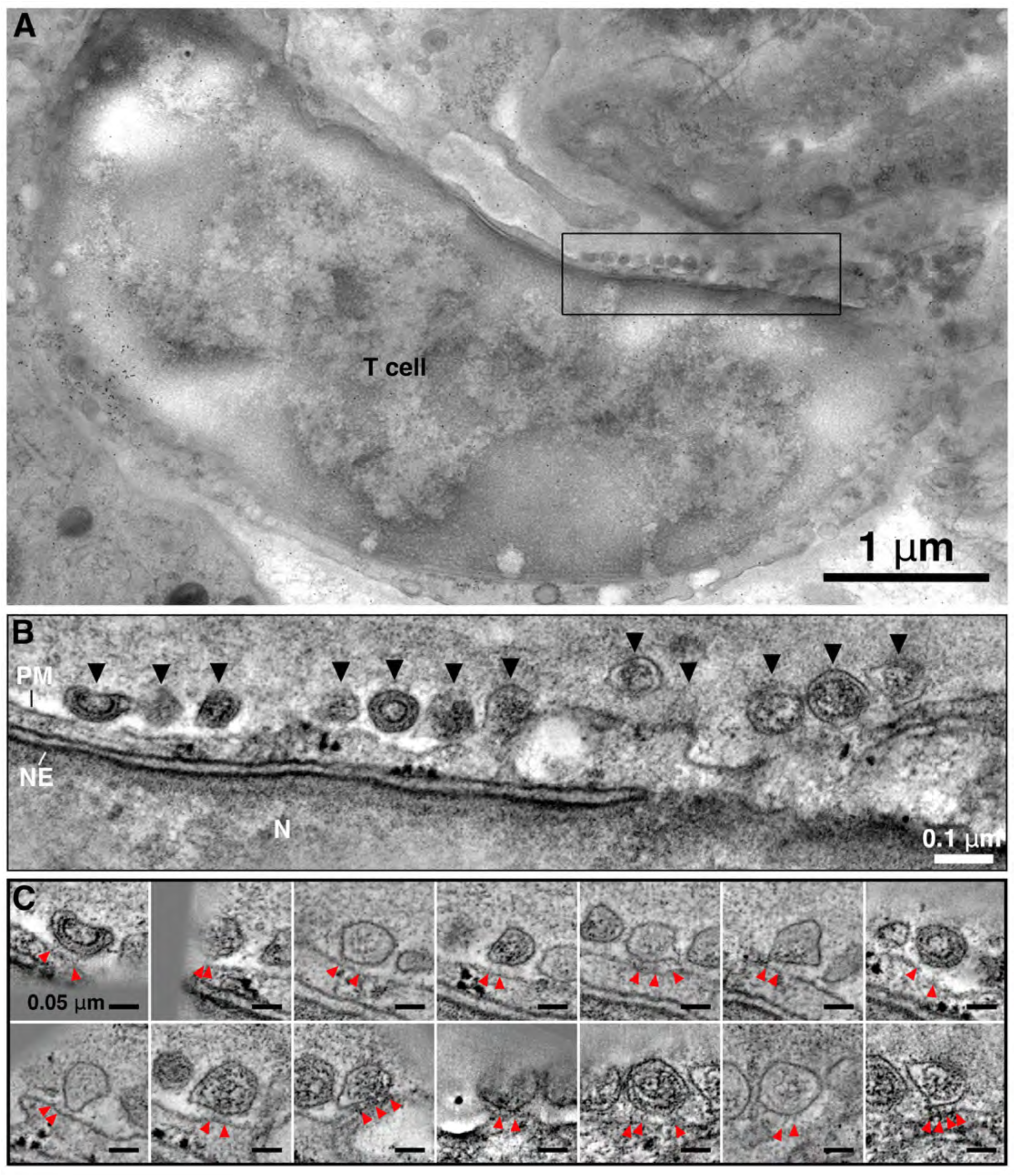
Trapped virions on the surface of a T cell in bone marrow. (A) T cell showing 14 HIV-1 virions trapped at the cell surface (black rectangle). (B) Tomographic reconstruction of the region indicated by the black rectangle in A, showing 12 of 14 trapped virions at the T cell surface (2 virions are outside of the plane of this tomographic slice) (PM, T cell plasma membrane; NE, T cell nuclear envelope; N, T cell nucleus). (C) Gallery of tomographic slices (1.6 nm) of each of the 14 fusing virions, rotated in 3D to optimize visualization of spokes (red arrowheads).

**FIG 5.**
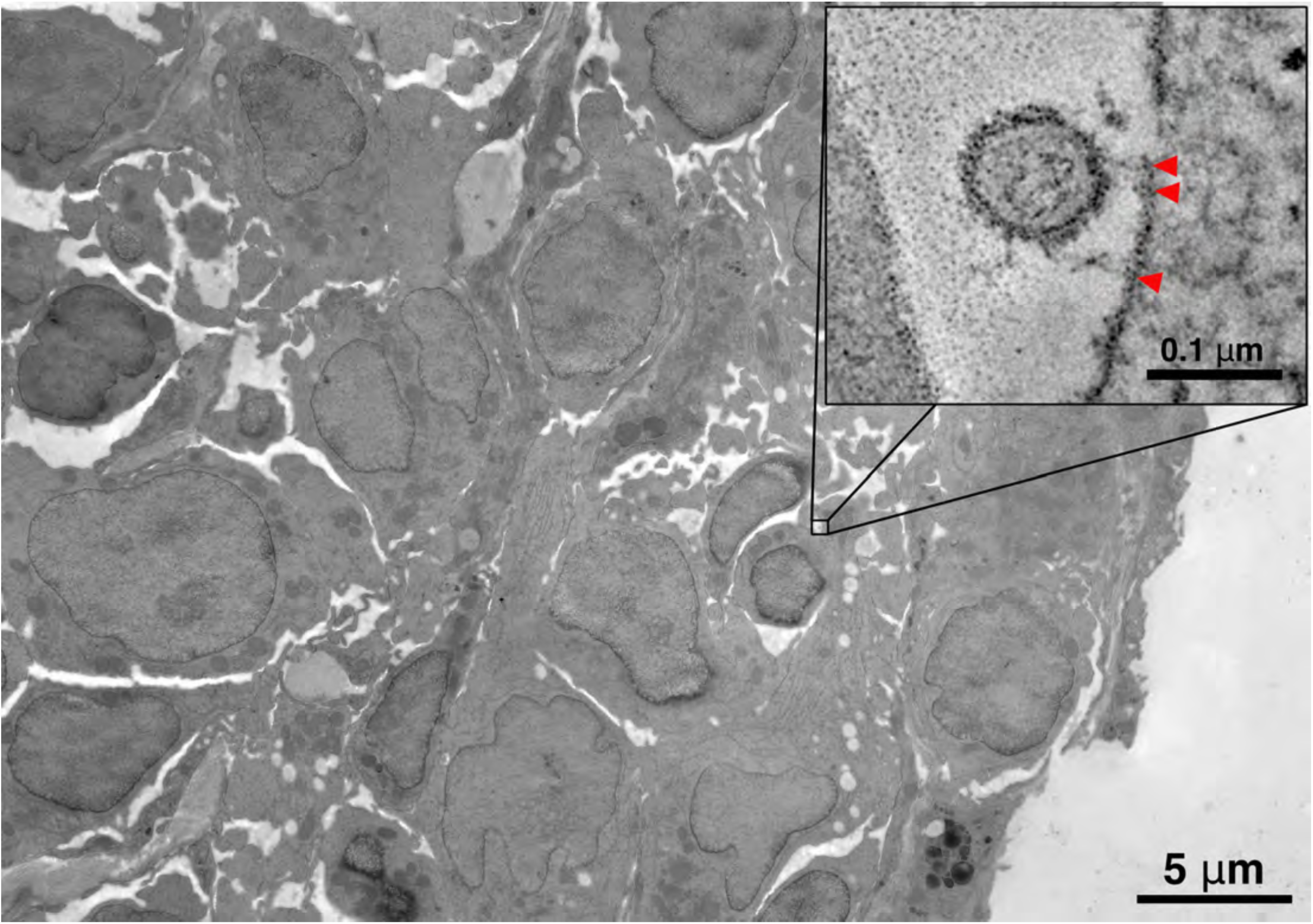
Trapped virions in the mesenteric lymph node. HIV-1 virions were found associated with immune cells, enmeshed among other cells comprising a germinal center. Inset: Tomography shows a mature HIV-1 virion trapped at the surface of a target cell by 3 spokes (red arrowheads).

**FIG 6.**
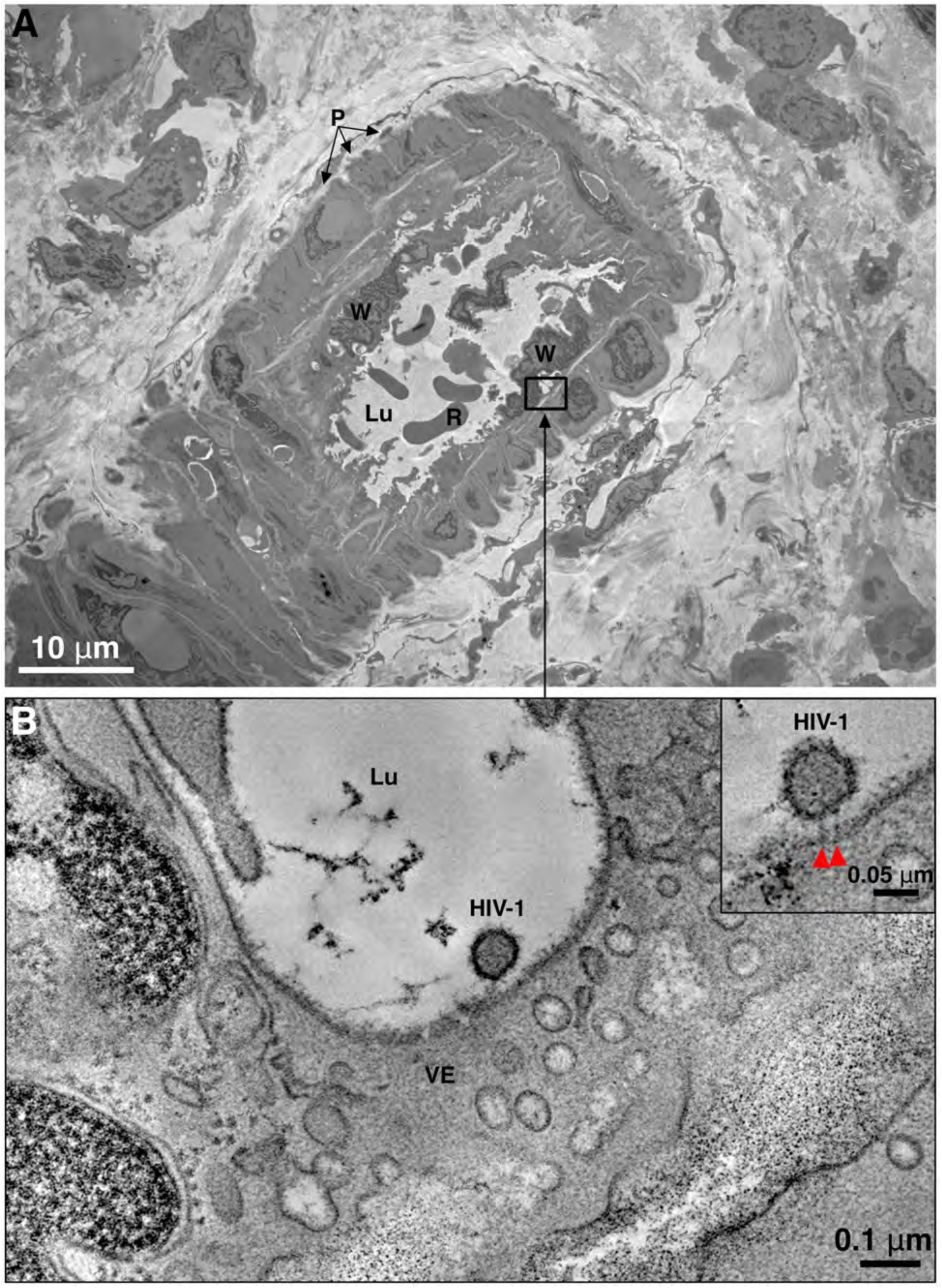
Trapped virions in mesenteric lymph node. (A) Montaged overview of a pericyte-laden blood vessel in mesenteric lymph node. P, pericytes; W, white blood cells, R, red blood cells, Lu, blood vessel lumen. (B) Tomographic slice (0.9 nm) of the region indicated by the square in A, showing a mature HIV-1 virion associated with the surface of a vascular endothelial cell (VE). Inset: Detail of the HIV-1 virion from the tomogram, rotated in 3D to reveal two spokes (red arrowheads), indicating that the virion is trapped at the surface of the endothelial cell.

**FIG 7.**
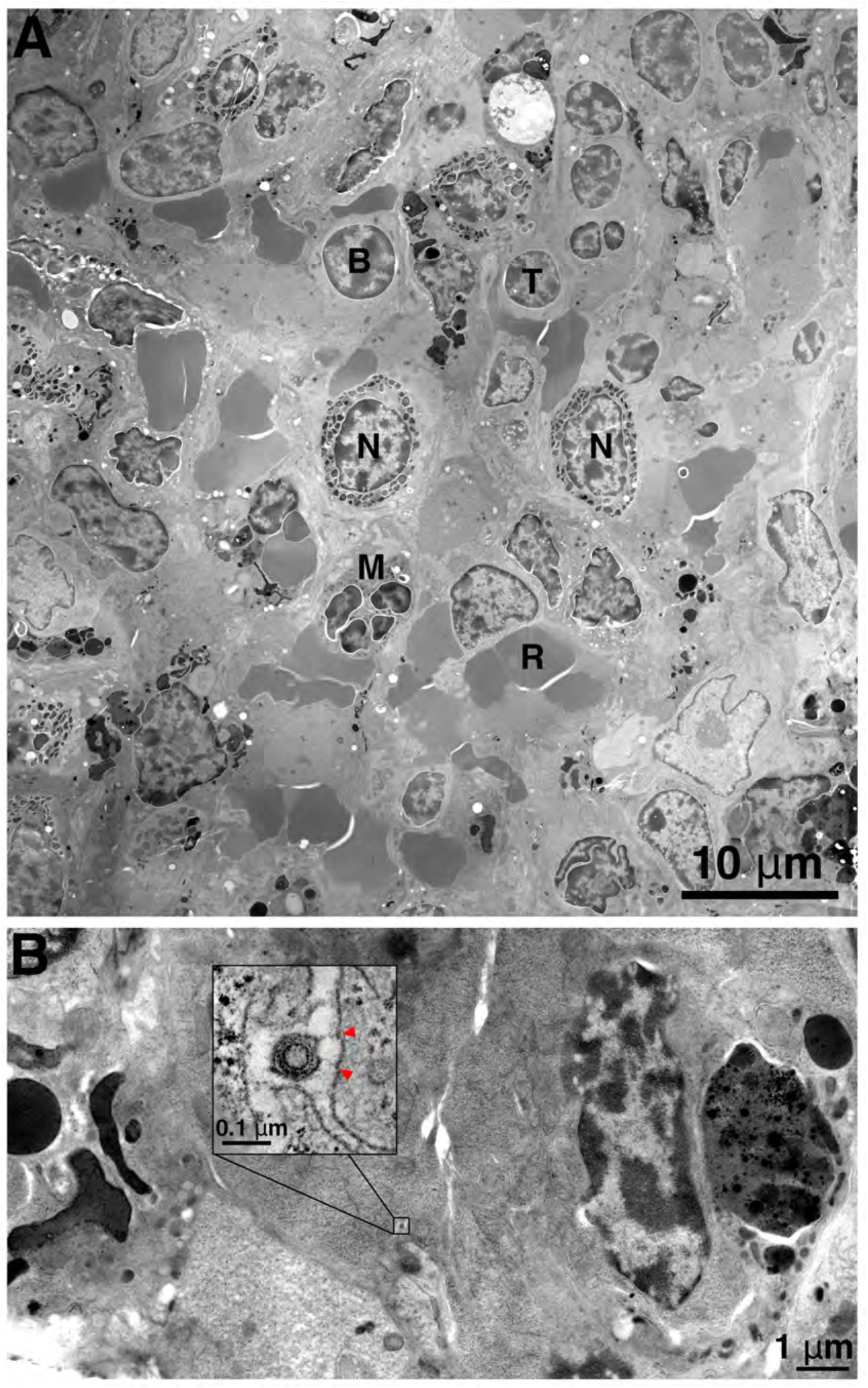
Trapped virions in spleen. (A) Overview of a region of white pulp in HIV-1–infected, CPT31-treated humanized BLT mouse spleen. Various types of immune cells, including neutrophils (N), polymorphonuclear macrophages (M), T cells (T) and B cells (B) are tightly apposed with other splenic cells and red blood cells (R). (B) Region of two closely apposed splenic cells containing an inhibitor-trapped immature HIV-1 virion. (Inset) Tomographic slice (0.9 nm) of the immature virion with two spokes (red arrowheads) attaching it to an adjacent cell surface, demonstrating CPT31 effectiveness in the spleen and identifying the attached cell as an HIV-1 target.

### Trapped HIV-1 virions in GALT

We first chose to search for CPT31-trapped virions in the GALT of HIV-1–infected humanized BLT mice to complement earlier ET studies (19, 21) and because GALT is a repository of large amounts of HIV-1 during acute infection (38, 39). Indeed, the GI tract contains a majority of CD4^+^/CCR5^+^ activated memory T cells that are targets of HIV-1 infection (40, 41), and T cells in GALT are among the first to be depleted as infection progresses from acute to chronic stages (42, 43).

GALT samples from the mouse small intestine were oriented to allow surveying of apical (epithelium) and basolateral (crypts of Lieberkühn and mucosa) regions. Similar to our previous ET study of HIV-1–infected humanized BLT mouse GALT (21), HIV-1 virions were primarily found in pools within intercellular spaces in the lower portions of crypts (Fig. 2, Fig. 3A-C). The numbers of virions in pools ranged from two to several dozen (Fig. 3C). By examining individual virions in varying 3D tomographic orientations, we resolved spokes linking virions to adjacent cell surfaces, confirming that the CPT31 inhibitor functions in GALT and allowing us to distinguish trapped from untrapped virions and identify HIV-1 target cells within the tissue (Fig. 2, Fig. 3D). The presence of nascent budding profiles, when found, further allowed us to identify HIV-1 producer cells within the same tissue regions (Fig. 1B). Trapped virions typically displayed 2-4 spokes linking them to the surface of target cells, similar to our previous findings in cultured cells (22). Three-dimensional analysis and segmentation of trapped virions (Fig. 3D) provided an assessment of spoke structure: measurement of 58 spokes from 21 GALT ET datasets showed an average spoke length of 18.4 ± 5.0 nm and spoke width of 3.6 ± 0.8 nm (Fig. 3E), consistent with estimates of pre-hairpin intermediate dimensions (Fig. 1A). The majority of trapped virions (∼80%) in GALT were mature, as distinguished by their one-shaped cores. Trapped immature virions identified by C-shaped cores (35, 36) were also found.

### Trapped HIV-1 virions in Bone Marrow

Bone marrow is the originating organ for many lymphoid cells, all of which are of human origin in humanized BLT mice due to transplantation of human hematopoietic stem cells (24). The relatively sparse distribution of cells in bone marrow facilitates locating and identifying specific HIV-1 target cells and any virions that may be associated with them. In our previous study (20), HIV-1 virions were pervasive within humanized BLT mouse bone marrow and observed as buds from infected T cells, transiting towards target cells, and associating extensively with macrophages. In macrophages, virions were found within phagocytic compartments along with remnants of previously phagocytosed cells and also budding into specialized cytoplasmic compartments that fused with invaginations of the plasma membrane to facilitate extracellular transmission (20).

Similar to our previous results (20), HIV-1 virions in CPT31-treated humanized BLT mouse bone marrow were found in proximity to lymphoid cells that could be identified as HIV-1 target cells: e.g., T cells were identified as small, round cells with comparatively large nuclei relative to their cytoplasmic volume; macrophages by their extensive surface invaginations and polymorphonuclear nature, and neutrophils by the presence of large, dense cytoplasmic granules. These criteria were used to identify lymphoid cells in other tissues in this study. In contrast to our previous studies (19–21), both mature and immature HIV-1 virions in tissues from CPT31-treated animals were often found in close contact with cell surfaces, and ET revealed that the majority of these virions exhibited spoke structures indicative of fusion inhibitor trapping (Fig. 4) with characteristics resembling trapped virions in GALT (Fig. 2, 3). In bone marrow, multiple HIV-1 virions were also observed in a line along the surface of a target cell (Fig. 4A,B). In such cases, 3D analysis of individual virions revealed spokes on each, indicating that all virions in the line were trapped (Fig. 4C). This contrasts with previous ET results in bone marrow from non-fusion inhibitor-treated samples (20), in which multiple virions were never observed in lines along the surface of a cell, suggesting that fusion inhibitors could slow HIV-1 ingress to a point at which large numbers of virions can be caught and imaged during the fusion process.

### Trapped HIV-1 virions in the Mesenteric Lymph Node

Lymph nodes are highly vascular by nature, allowing continuous circulation of lymphoid cells from the bloodstream and lymphatic system (44). Thus, we can look in infected humanized BLT mice for HIV-1 in follicular regions, germinal centers, and within the blood and lymph vessels that intercalate through the organ. Cells comprising the lymph node germinal center are tightly packed together, but lymphoid cells (potential HIV-1 targets) were identified by criteria described above for bone marrow samples. When located, the surfaces of immune cells and the extracellular spaces between them were scanned for the presence of HIV-1. Virions were identified by the same structural criteria used for other tissues in this study, and spokes were characterized by the same 3D tomographic analyses. As expected, individual HIV-1 virions were found associated with T cells in germinal centers (Fig. 5), and their pre-hairpin intermediate trapped status could be determined by observing spokes by ET (inset, Fig. 5).

In addition, HIV-1 virions were found within microvascular blood vessels, associating with endothelial cells and with peripheral pericytes (Fig. 6). Endothelial cells comprise the inner lining, or wall, of the vessel, and pericytes wrap around the endothelial cells of capillaries and venules. Both endothelial cells and pericytes express CD4 and HIV-1 co-receptors (45), characterizing them as HIV-1 target cells (46–49). In the case of pericytes, HIV-1 infection of blood–brain barrier pericytes has been found in post-mortem samples of human brains and in a mouse model of HIV-1 infection (45, 50). Our finding of HIV-1 virions trapped to the surfaces of vascular endothelial cells and pericytes (inset, Fig. 6) confirms that the CPT31 inhibitor functions in lymph nodes of HIV-1–infected humanized BLT mice and reaffirms that fusion inhibitors can be used as markers for HIV-1 target cells within tissues of an infected animal.

### Trapped HIV-1 Virions in Spleen

The spleen is a multifunctional organ that is highly enriched in lymphoid cells. Approximately 80% of splenic tissue is red pulp, where blood is filtered to remove damaged or dead cells and larger contaminants. The remaining 20% is white pulp, which comprises T cells, B cells, and macrophages and thus plays a critical role in immune responses (51).

When dissecting splenic tissue, white pulp can be distinguished from red pulp at the histological level and specifically selected for EM preparation. Although cells are tightly apposed, lymphoid cells could be identified and characterized by the same criteria used for lymphoid cells in bone marrow and other tissues in this study. We found that HIV-1 virions were fewer in number in spleen than in other tissue samples but could be found by scanning cellular peripheries and intercellular spaces. Once found, trapped virions were identified by the presence of 2-4 spokes per virion attaching them to cells. The attached cells were thereby characterized as HIV-1 targets (Fig. 7). Trapped mature versus immature HIV-1 virions were found in roughly similar proportions within the spleen.

## DISCUSSION

We expanded upon our previous ET study of fusion inhibitor-treated HIV-1 pseudoviruses and cultured host cells (22) to examine optimally preserved HIV-1–infected tissues from humanized mice treated with the potent fusion inhibitor CPT31 (12). The goal of both studies was to visualize virions trapped by pre-hairpin intermediate structures in which the gp41 transmembrane region and the fusion peptide link the viral and target cell membranes (8). Trapped HIV-1 virions were found in all tissues that we examined (small intestine, mesenteric lymph nodes, spleen, and bone marrow) and could be confidently distinguished from untrapped virions in the same tissue regions. Using ET and 3D reconstructions, presumptive pre-hairpin intermediates linking virions to host cell surfaces were visualized as 2-4 narrow spokes per trapped virion. The spokes linking virions to cell surfaces were similar in number and structure to those seen in our previous study in cultured cells (22). Based on their dimensions, the spokes likely correspond to extended Env trimers after gp120 dissociation but prior to collapse into the post-fusion six-helical bundle structure (8). Our observation of relatively few (2–4) spokes per attached virion in both this study and our previous in vitro ET study (22) implies that HIV-1 Env-mediated membrane fusion requires only a fraction of the ∼14 Envs on each virion (52–56).

In all tissue samples examined for this study, we found that the majority (∼80%) of trapped virions were mature, as demonstrated by conical cores. Immature HIV-1 virions are not infectious until after proteolytic processing of Gag and Gag-pol proteins, which also frees the viral capsid to adopt a cone shape (57). We also found ∼20% of trapped virions in tissues were immature, as demonstrated by the presence of a “C-shaped” nascent viral core. This is consistent with immature HIV-1 virions presenting Env trimers that can initiate fusion with a target cell. However, given that immature HIV-1 particles are up to 10-fold less active for fusion than mature HIV-1 (58), we may be observing a larger than representative percentage of immature virions trapped by the CPT31 fusion inhibitor due to presumably slower fusion kinetics for immature virions.

The fusion inhibitor used in this study, CPT31, is a promising HIV-1 antiviral, having been shown to act as a potent inhibitor of HIV-1 entry and exhibit a favorable pharmacokinetic profile and efficacy as a preventative and therapeutic agent in a non-human primate model of HIV-1 infection (12, 59, 60). Of relevance to potential therapeutic use of CPT31, the finding of trapped HIV-1 virions in all tissues examined for this study demonstrates a wide distribution of the CPT31 inhibitor within a treated animal. In addition, the presence of virions trapped by spokes, particularly in vascular endothelial cells, demonstrates that the CPT31 fusion inhibitor can be used as a marker for potential HIV-1-target cells within tissues, facilitating the mapping of HIV-1 target cells within the complex cellular milieu of infected tissues.

## ACKNOWLEDGEMENTS

The authors thank Brett Welch for providing CPT31, Michael Brehm and Dale Greiner (U Mass Chan Medical School) for providing BLT mice, the Caltech Cryo-EM Center for maintenance of the T12 electron microscope, Walther Mothes and members of the Bjorkman and Kay laboratories for helpful discussions and critical review of the manuscript. This work was supported by NIH U54 AI170856 (MSK and PJB), NIH R01AI145164 (PK), and CIHR grant #422148 (PK).

## MATERIALS AND METHODS

### CPT-31 inhibitor preparation

CPT31 was synthesized and purified as described (12). Stock solutions were prepared at 2.5 mg/mL in PBS.

### HIV-1 infection of humanized BLT mice and inhibitor treatment regimen

Humanized BLT mice (NSG-BLT) were generated as previously described using immunodeficient NOD.Cg-*Prkdc^scid^Il2rg^tm/Wjl^*/Sz (NOD-*scid IL2rγ^null^*) mice from the Jackson Laboratory (Bar Harbor, ME) (61, 62) and tested for engraftment at 10 weeks after transplant for levels of human hematopoietic cells by multicolor flow cytometric analysis using mAbs: anti-human CD3, CD4, CD8, CD11c, CD19, and CD45; anti-mouse CD45; and isotype control mAbs (Biolegend, Inc.).

Mice were intraperitoneally infected with HIV-1_JRCSF_ for 3 weeks prior to treatment with inhibitor at which time plasma viral loads were >10^5^ copies/mL. Mice received two doses of CPT31 inhibitor (10 mg/kg/dose), IV-retro-orbital, 2 h apart. Mice were euthanized 2 h following the second dose and tissue samples collected. This dose will provide a plasma drug concentration >5 µM at the time of sample collection, >1000-fold CPT31’s estimated IC_50_ based on data in ref. (12).

Tissue samples, including intestine (GALT), spleen, mesenteric lymph node and bone marrow, were removed from the animal and immediately prefixed with 3% glutaraldehyde, 1% paraformaldehyde, 5% sucrose in 0.1 M sodium cacodylate trihydrate at 4°C. For bone marrow samples, a hind leg was removed and the marrow extracted from the femur bone in subsequent processing steps.

### Sample preparation for electron microscopy

Tissue samples were rinsed with 0.1 M cacodylate buffer and dissected into suitable-sized pieces, rinsed with fresh cacodylate buffer containing 10% Ficoll, placed into brass planchettes (Ted Pella, Inc.), and rapidly frozen with an HPM-010 high-pressure freezing machine (Bal-Tec, Lichtenstein). The frozen samples were transferred under liquid nitrogen to cryotubes (Nunc) containing a frozen solution of 2.5% osmium tetroxide, 0.05% uranyl acetate in acetone. Tubes were loaded into an AFS-2 freeze-substitution machine (Leica Microsystems) and processed at −90°C for 72 h, warmed over 12 h to −20°C, held at that temperature for 6 h, and then warmed to 4°C for 2 h. The fixative was removed, and the samples rinsed 4x with cold acetone, after which they were infiltrated with Epon-Araldite resin (Electron Microscopy Sciences) over 48 h. The tissue samples were flat-embedded between two Teflon-coated glass microscope slides and the resin was polymerized at 60°C for 48 h.

### Electron microscopy and dual-axis tomography

Embedded tissues were observed by light microscopy and appropriate regions were extracted with a microsurgical scalpel and glued to the tips of plastic sectioning stubs. Semi-thin (170 nm) serial sections were cut with a UC6 ultramicrotome (Leica Microsystems) using a diamond knife (Diatome, Ltd). Sections were placed on formvar-coated copper-rhodium slot grids (Electron Microscopy Sciences) and stained with 3% uranyl acetate and lead citrate. Gold beads (10 nm) were placed on both surfaces of the grid to serve as fiducial markers for subsequent image alignment. Grids were placed in a dual-axis tomography holder (Model 2040, E.A. Fischione Instruments) and imaged with a Tecnai T12-G2 transmission electron microscope operating at 120 KeV (ThermoFisher Scientific) equipped with a 2k x 2k CCD camera (XP1000; Gatan, Inc.). Tomographic tilt-series and large-area montaged overviews were acquired automatically using the SerialEM software package (63). For tomography, samples were tilted ± 62° and images collected at 1° intervals. The grid was then rotated 90° and a similar series taken about the orthogonal axis. Tomographic data was calculated, analyzed, and modeled using the IMOD software package (64–66) on iMac Pro and Mac Studio M1 computers (Apple, Inc.).

---

**Movie 1.**
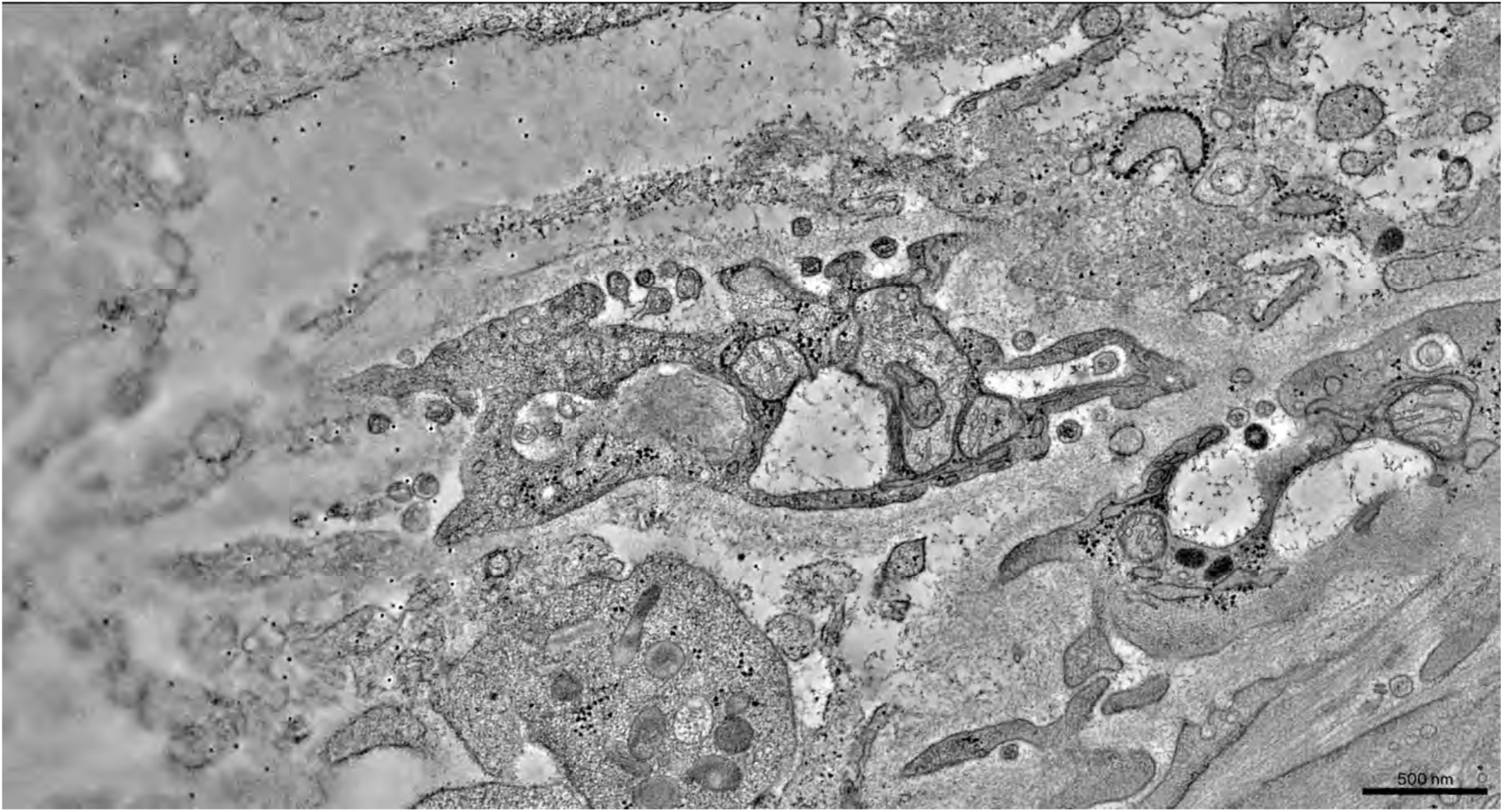
Identification and segmentation of inhibitor-trapped virions in HIV-1–infected, CPT31-treated humanized BLT mouse bone marrow. Montaged tomographic reconstruction of a macrophage in bone marrow, associated with mature and immature HIV-1 virions. The movie shows the volume of the reconstruction (170 nm) and then increases the magnification and pans across the volume along the y-axis, further zooming in to a specific mature virion. Viewing the virion through its volume reveals three spokes linking it to the adjacent cell’s surface, further illustrated by segmentation (green, virus periphery; blue, cone-shaped core; magenta, spokes) and rotation of the model in 3D.

